# Comparing genome-wide association study results from different measurements of an underlying phenotype

**DOI:** 10.1101/313445

**Authors:** Joseph L. Gage, de Leon Natalia, Clayton Murray

## Abstract

Increasing popularity of high-throughput phenotyping technologies, such as image-based phenotyping, offer novel ways for quantifying plant growth and morphology. These new methods can be more or less accurate and precise than traditional, manual measurements. Many large-scale phenotyping efforts are conducted to enable genome-wide association studies (GWAS), but it is unclear exactly how alternative methods of phenotyping will affect GWAS results. In this study we simulate phenotypes that are controlled by the same set of causal loci but have differing heritability, similar to two different measurements of the same morphological character. We then perform GWAS with the simulated traits and create receiver operating characteristic (ROC) curves from the results. The areas under the ROC curves (AUCs) provide a metric that allows direct comparisons of GWAS results from different simulated traits. We use this framework to evaluate the effects of heritability and the number of causative loci on the AUCs of simulated traits; we also test the differences between AUCs of traits with differing heritability. We find that both increasing the number of causative loci and decreasing the heritability reduce a trait’s AUC. We also find that when two traits are controlled by a greater number of causative loci, they are more likely to have significantly different AUCs as the difference between their heritabilities increases. These results provide a framework for deciding between competing phenotyping strategies when the ultimate goal is to generate and use phenotype-genotype associations from GWAS.

## Introduction

As image-based methods for quantifying plant phenotypes grow in popularity, they present the ability to measure phenotypes that previously could not be easily quantified as well as an alternative way to measure phenotypes that previously had to be manually quantified. The types of novel phenotypes enabled by image analysis include fractal dimension (Gage *et al.* 2017), principal component analysis of plant organ biomass (Miller *et al.* 2016) or shape (Chitwood *et al.* 2014), and topological methods for quantifying branching patterns (Li *et al.* 2017). Image-based phenotyping enables increases in resolution, such as in Miller *et al.* (2016) scanning maize ears on a flatbed scanner at 1,200 dots per inch; it enables advances in accuracy, such as quantifying disease resistance (Bock *et al.* 2010); and it enables scalability and throughput, by building multiple imaging devices (Durham Brooks *et al.* 2010) or using a mobile imaging device (Men *et al.* 2012). However, image-based phenotype measurements are not always as accurate as high quality manually measured phenotypes, though the decrease in accuracy may be paired with an increase in throughput or decrease in cost if high quality manual phenotypes are time consuming or expensive to measure. Though it is not strictly an image-based method of phenotyping, one example of the tradeoff of accuracy for efficiency is the use of near infrared reflectance spectroscopy (NIRS), which has been used for decades as a way to predict chemical composition of silage feedstock without costly, expensive, and sometimes hazardous wet lab assays (Park *et al.* 1998).

Potential tradeoffs between measurement accuracy and throughput need to be carefully considered by scientists preparing for large-scale experiments. In the fields of plant breeding and plant genetics, genetic mapping experiments are one example of the type of study where such considerations are crucial. Genome-wide association studies (GWAS) involve measuring a phenotype, usually in a replicated population of hundreds to thousands of genetically distinct individuals, then scanning for statistical associations between individuals’ phenotype and their genotype at numerous genetic loci. In such studies, the accuracy and precision with which a phenotype is measured will have a direct impact on the ability to detect genetic associations by GWAS.

At its core, GWAS involves testing for a difference in phenotype between individuals with different genotypes at a particular single nucleotide polymorphism (SNP). This process is repeated separately for hundreds of thousands of SNPs across the genome. Ideally, SNPs within or near genes that have some effect on the phenotype of interest will result in strong statistical associations. As such, the precision and accuracy of phenotypic measurements influence the ability to detect statistical differences between groups of individuals with different alleles. The heritability of a phenotype, defined as the ratio of genotypic variance to phenotypic variance, is a useful way to quantify the proportion of phenotypic variability that is attributable to genetic differences between individuals. All other components held equal, heritability will increase as precision of phenotypic measurement increases, due to decreasing phenotypic variability from measurement error. Two methods of measuring the same ‘true’ phenotype with differing precision will have different heritability, and thus different power to detect SNPs that are statistically associated with the ‘true’ phenotype of interest. For the remainder of this study we will refer to the ‘true’ phenotype of an individual as its character, and will refer to different measurements of a character as traits.

In addition to heritability, another parameter that affects power in GWAS is the number of loci that control a character. For two characters measured with the same heritability, one controlled by fewer loci will have on average a larger proportion of variance explained by each locus. Power to detect an association at a particular locus is positively related to the proportion of phenotypic variance explained by that locus (Visscher 2008). Thus, phenotype-genotype associations for characters controlled by a large number of small-effect loci tend to be more difficult to detect.

Increased throughput of image-based phenotyping methods can make it possible to collect measurements of more traits on more individuals than by manual measurement, which makes image-based phenotyping an attractive way to generate phenotypic data for GWAS. We can consider the manual measurement and the image-based measurement of an individual character to be two traits with differing heritability but the same exact set of causative loci. As in the NIRS example above, researchers may sometimes prefer a less precise method for measuring a character because it is cheaper, faster, or otherwise more desirable. It is unclear just how much loss in precision (decrease in heritability) can occur before GWAS results begin to suffer. Part of the answer to this question lies in the goals and risk tolerance of the researcher: if the goal of the experiment requires identification of few, strong signals then perhaps lower power to detect associations will still be tolerable; if instead the goal is to identify many small-effect loci, then even small reductions in heritability could negatively impact the outcome of the study. In this experiment, we use simulations to investigate the relationship between trait heritability and the ability to detect genetic regions associated with a character. We use receiver operating characteristic (ROC) curves to characterize detection of causative loci. Previous studies have used ROC curves or similar visual aids to evaluate the efficacy of different GWAS methods (e.g., Wang *et al.* 2014; Liu *et al.* 2016). In this study, however, we use ROC curves to evaluate GWAS results for simulated traits, and test for differences between those ROC curves using the area under the curve (AUC). Using AUC to distill GWAS results to a single statistic enables direct comparison of GWAS results from traits with differing simulation parameters. We use a test statistic for differences between two AUCs to construct a null distribution from simulated traits with the same parameters, and use that distribution to predict whether real traits measured manually and by image analysis have significantly different AUCs. These results provide a framework for evaluating how differences in heritability between two measures of a character can impact the efficacy of GWAS for identifying loci associated with the character of interest.

## Results

### Manual and image-based phenotypes are highly correlated

Empirical phenotypic data used in this study comes from morphological measurements of the male inflorescence of maize, known as the tassel. Four tassel morphological characters were measured manually and by image analysis in the Wisconsin Diversity panel, a population of 942 diverse inbred maize lines (WiDiv-942). The manually measured traits were tassel length (TL), spike length (SL), branch number (BN), and tassel weight (TW), and their image-based counterparts are referred to as TLp, SLp, BNp, and TWp, respectively. TL, SL, and BN were measured in replicated experiments over three years, while the other five traits were measured in a replicated experiment in one year. Best linear unbiased predictors (BLUPs) for manually measured traits are highly correlated with BLUPs for the corresponding image-based traits, with Pearson’s correlation coefficients ranging from 0.81 to 0.9 (Figure 1) (Gage *et al.* 2018). Estimated heritability for the traits ranges from 0.95 to 0.97 for manually measured traits and from 0.79 to 0.86 for image-based traits (Table 1) (Gage *et al.* 2018). TL, SL, and BN were measured in three environments, and all image-based traits as well as TW were measured in a single environment. The difference in number of environments could inflate the heritability estimates of the image-based traits, making it reasonable to conclude that the image-based heritability estimates represent an upper bound for their true heritabilities. As such, it is reasonable to conclude that image-based measurements are less precise than the manual measurements.

**Figure 1:**
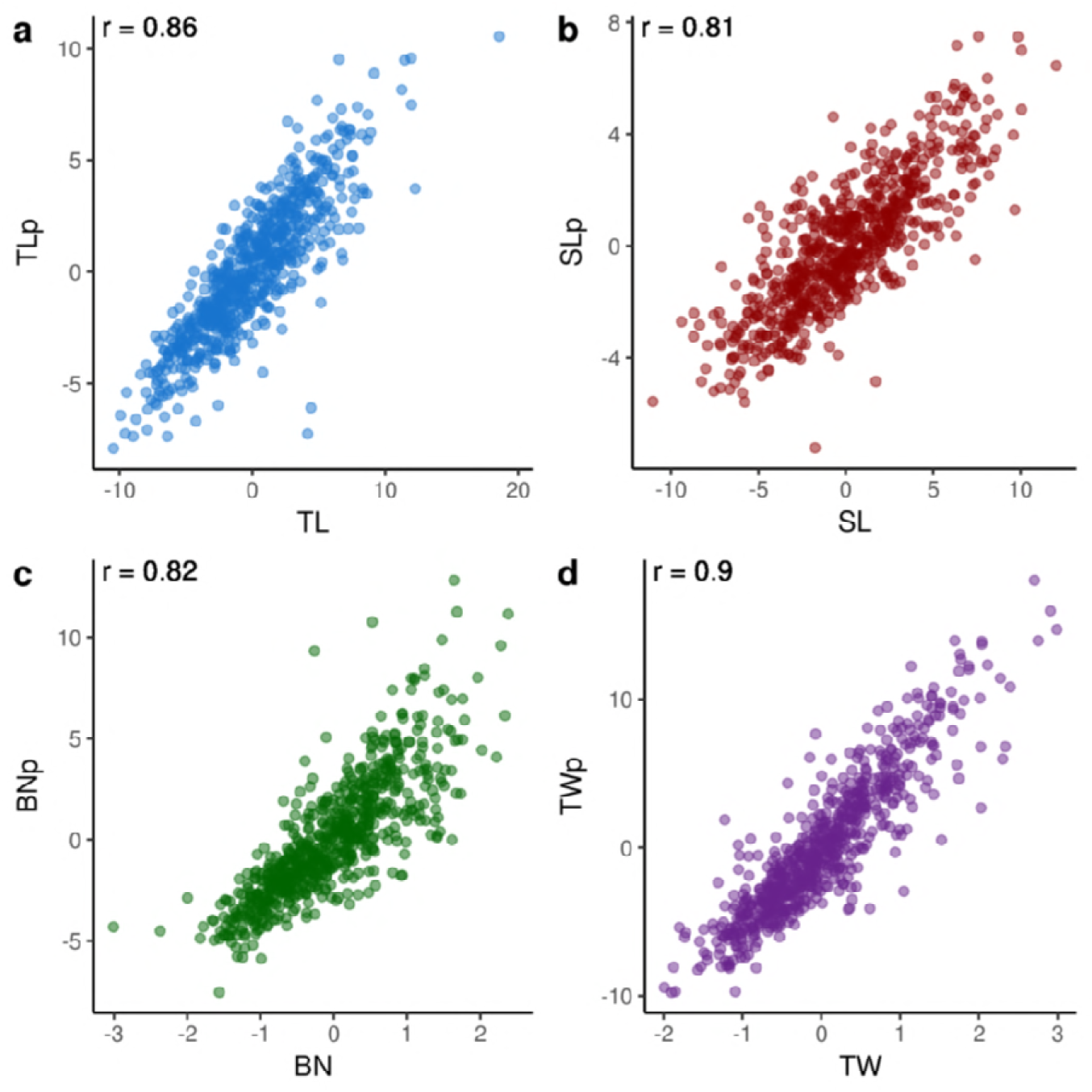
Correlations between manual and image-based phenotypic values. Scatter plots of best linear unbiased predictors (BLUPs) for manual vs image-based measurements of tassel length (TL; a), spike length (SL; b), branch number (BN; c), and tassel weight (TW; d). Manually measured BLUPs are along the x-axis, while image-based measurements are on the y-axis. Values in the upper left corner of each plot are the Pearson correlation coefficients for each trait.

**Table 1:**
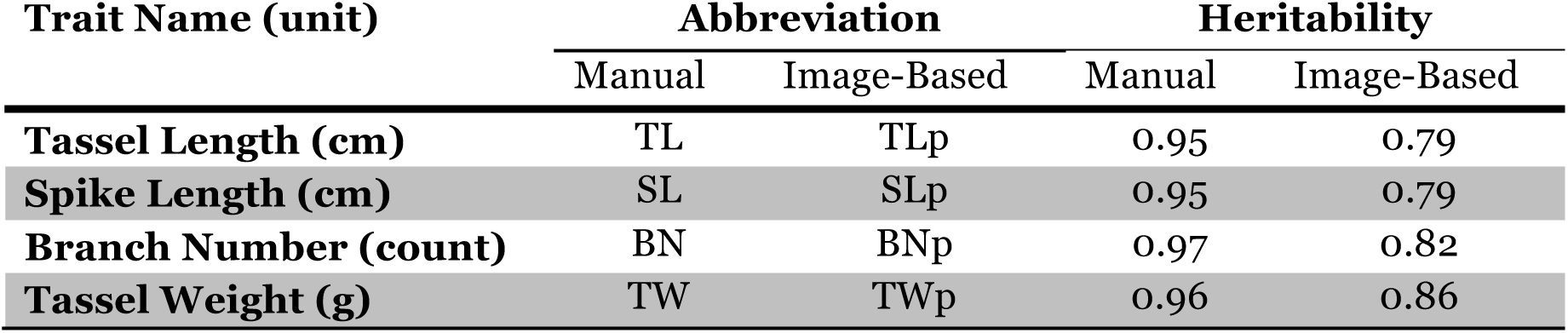
Comparison of manual and image-based trait heritabilities. Heritabilities for four different tassel morphological traits, measured both manually and using image-based methods. TL, SL, and BN were measured in three environments, whereas TW, TLp, SLp, BNp, and TWp were measured in one environment.

| Trait Name (unit) | Abbreviation |  | Heritability |  |
| --- | --- | --- | --- | --- |
|  | Manual | Image-Based | Manual | Image-Based |
| **Tassel Length (cm)** | TL | TLp | 0.95 | 0.79 |
| **Spike Length (cm)** | SL | SLp | 0.95 | 0.79 |
| **Branch Number (count)** | BN | BNp | 0.97 | 0.82 |
| **Tassel Weight (g)** | TW | TWp | 0.96 | 0.86 |

### Power in GWAS of simulated traits varies with heritability and number of causal loci

To create a framework for comparing traits measured manually and by image analysis, we first performed simulations to examine the impact of heritability and number of causative loci (NCL) on GWAS results. We first simulated a number of traits with different heritability and NCL. Phenotypes were simulated as being controlled by single nucleotide polymorphisms (SNPs) from the WiDiv-942, which was genotyped at 529,018 SNPs discovered by RNA sequencing. Simulated phenotypes were controlled by varying NCL: 10, 100, or 1,000 randomly selected SNPs were randomly assigned effect sizes drawn from a normal distribution. For each of the three values for NCL, traits were simulated with heritabilities ranging from 0.1 to 0.9 in increments of 0.1. Each combination of NCL and heritability was simulated ten times. GWAS were performed on all simulated phenotypes, and empirical ROC curves were created with the results from each GWAS. ROC curves plot the proportion of true positives (true positive rate; TPR) against the proportion of false positives (false positive rate; FPR), as the threshold for labeling an observation as positive moves from stringent to more liberal. The ROC curve for a test with very good ability to identify true positives without too many false positives will rise steeply from the origin and approach the point (0, 1), before flattening out and continuing on to the point (1,1), producing an AUC close to 1. A test that is no better than randomly guessing which observations are positives will yield an ROC curve that follows a line with slope equal to one from the origin to (1,1), producing an AUC of 0.5.

ROC curves are typically constructed by classifying a number of individuals as either cases or controls, based on some continuous predictor variable. The TPR and FPR are calculated at different levels of the predictor variable to create the curve. To create ROC curves for GWAS results each SNP is treated as an individual, the true status of which is either causative (case) or non-causative (control). The continuous predictor variable is the −log_10_(p-value) for each SNP from GWAS.

Our empirical results show that for any given NCL, simulated traits with higher heritability generally had better ROC curves, as measured by AUC (Figure 2). This was expected, as greater heritability implies greater genetic variance relative to error, which makes it easier to identify associations between phenotypic values and genotypic groups at causal loci. However, higher heritability does not guarantee better ROC curves as there are ROC curves with different heritability that intersect, particularly when the NCL is low (Figure 2).

**Figure 2:**
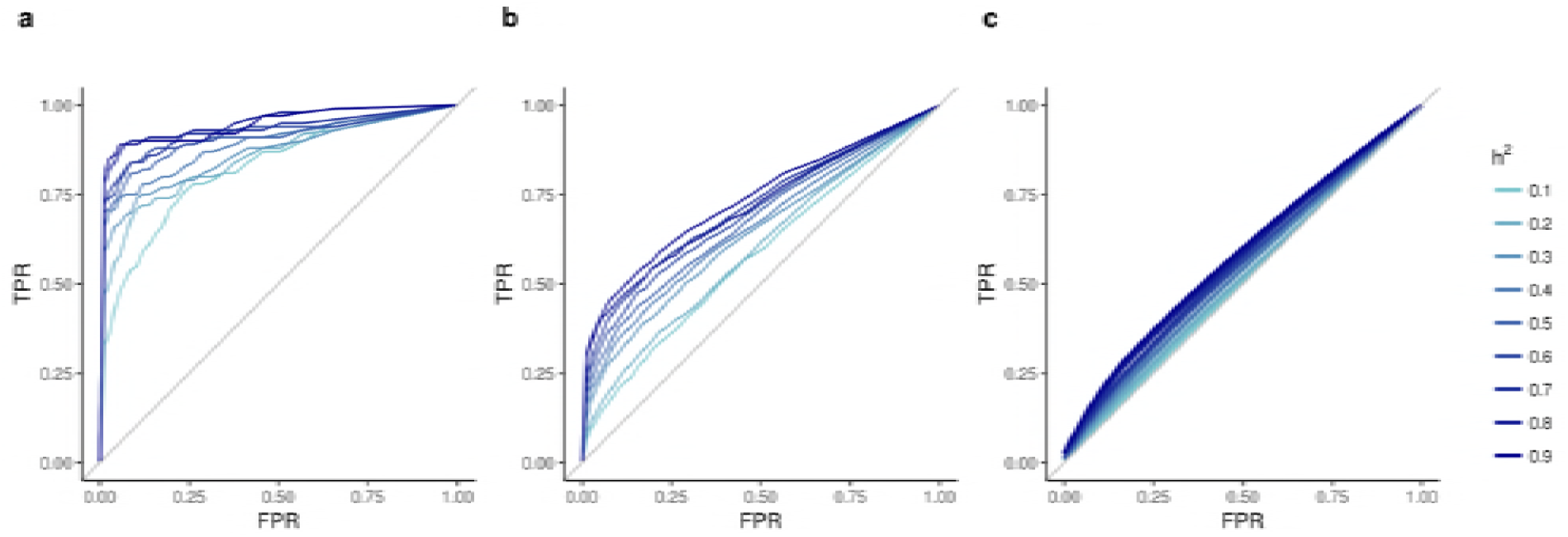
Receiver operating characteristic curves for GWAS of simulated traits. Receiver operating characteristic curves for GWAS results of traits controlled by 10 (a), 100 (b), or 1000 (c) causal loci. For each number of causal loci, simulation of traits with heritabilities ranging from 0.1 to 0.9 were replicated 10 times each. Each curve represents the average of the ten replications for each combination of causal loci and heritability. TPR: true positive rate; FPR: false positive rate; h^2^: heritability.

For any given heritability, the NCL plays an even larger role in the shape of the ROC curve, with traits controlled by more loci having worse ROC curves than those with fewer loci (Figure 2). This result was also expected. The effect sizes of individual loci become smaller as NCL increases, making detection of true associations more difficult.

### Heritability and number of causal loci influence ability to detect differences between ROC curves

The AUC can be interpreted as the probability that, given a randomly selected pair of one causative and one non-causative SNP, the predictor variable [-log_10_(p-value)] for the causative SNP will be greater than or equal to the predictor variable for the non-causative SNP (DeLong *et al.* 1988). AUCs provide a way to describe an ROC curve with a single value, and can be used for testing differences between ROC curves. The empirical AUC of an ROC curve is equivalent to the statistic generated by a Mann-Whitney test on the predictor scores of the cases and controls, or in the context of this study, the causative and non-causative SNPs. Therefore, treating the AUCs of two empirical ROC curves as Mann-Whitney statistics permits non-parametric testing of the difference between two AUCs, the test statistic of which (Z) follows a standard normal distribution (DeLong *et al.* 1988). However, because we cannot be certain that the GWAS results from this study satisfy the assumptions of a Mann-Whitney test or the assumptions for testing the difference between AUCs, we still use the Z statistic but do not make the assumption that the distribution of Z is normal. We assume that ROC curves created from GWAS results on traits with the same parameters (i.e., NCL and heritability) should not be significantly different from each other. Therefore, the empirical distribution of Z when the difference in heritability (D) between two traits equals zero is the distribution of Z under the null hypothesis that the AUCs of two ROC curves are the same.

We hypothesized that as D between two traits increased so would the Z score, corresponding to a difference between AUCs of the two traits. We tested all 90 AUCs with the same NCL (10 replications times nine levels of heritability) against each other, resulting in 4,005 test statistics for each NCL (comparisons were only made in one direction). The goal of this study is to assess how differing heritability of two measurements of the same underlying character affects GWAS results. Because manual and image-based measurements of a character have equivalent underlying genetic structure, we limited our comparisons to AUCs of simulated traits with the same NCL.

The Z scores for each pairwise test of two traits were regressed against D (Figure 3). As expected, the Z values get more extreme as D gets larger – this is a reflection of higher heritability traits generally having larger AUCs than lower heritability traits. Note that the tests were always done in a consistent direction; therefore we mostly observed results with positive Z scores. The relationship between D and Z is more extreme for simulated traits with greater NCL. Practically, this indicates that within the assumptions of these simulations, heritability plays a smaller role in the ability to detect GWAS associations when the trait is controlled by a small NCL. ROC curves for traits with more complex genetic architectures, however, deteriorate more quickly as heritability declines.

**Figure 3.**
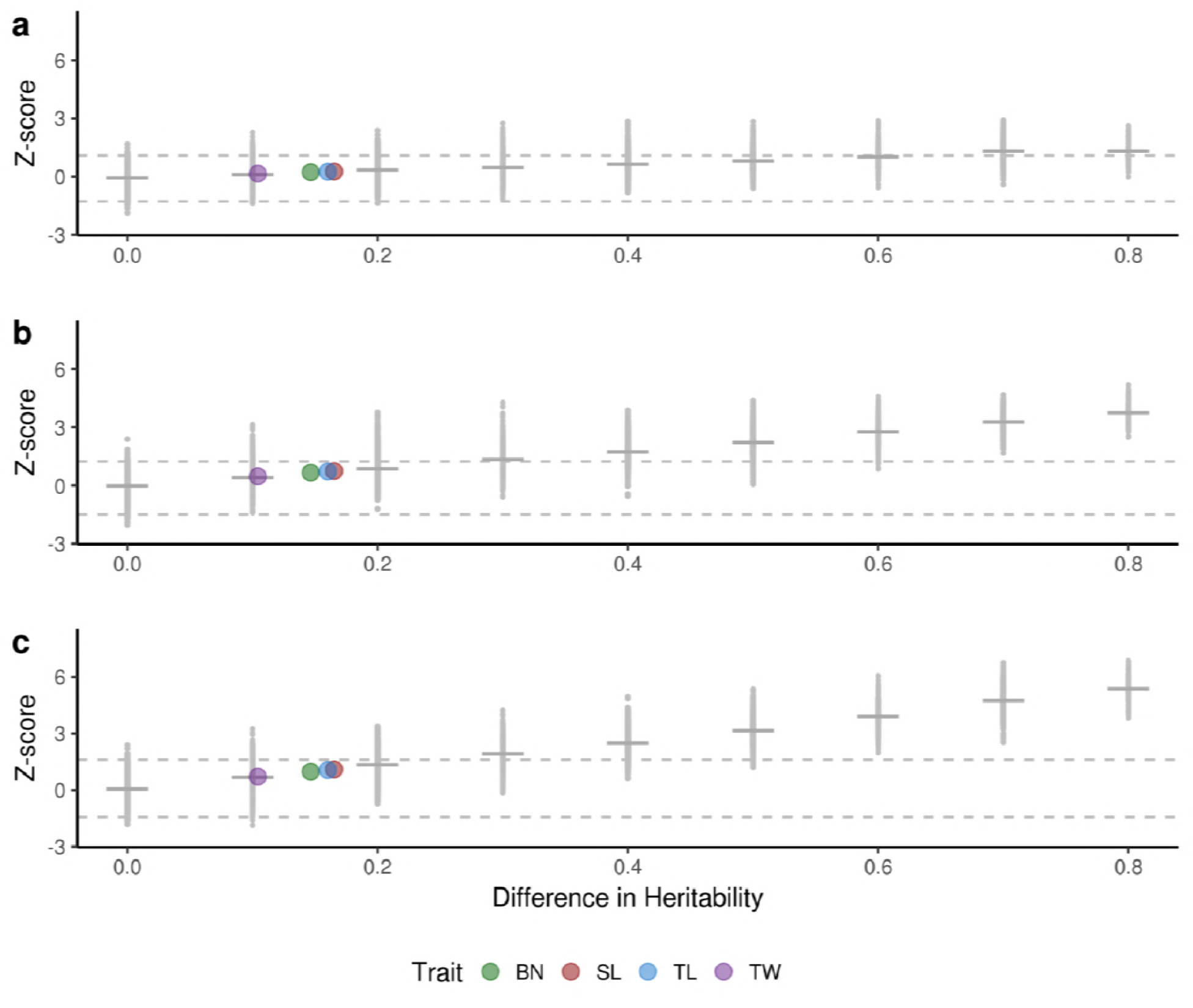
Results of testing area under the curve for simulated phenotypes. The Z score for testing the difference of two AUCs plotted against the absolute difference in heritability (D) between the two traits. Small gray dots represent the Z score from a single pairwise test between simulated traits, while horizontal gray bars represent the median Z score for a given D. Larger colored dots represent the D estimates for real traits, plotted along the line that best fits the Z scores of the simulated data. Dashed lines represent the thresholds for significance at α=0.05 (i.e., 2.5^th^ and 97.5^th^ percentiles), calculated from the empirical distribution of Z when D=0. BN: branch number; SL: spike length; TL: tassel length; TW: tassel weight.

### Alternative measurements of real phenotypes are not predicted to have differing AUCs

Having established an empirical relationship between Z and D for different NCL, we then used the results from our simulations to predict whether there is a significant difference between the AUCs of manual and image-based traits of a real character. We used the estimated heritabilities of manual and image-based measurements of TL, SL, BN, and TW to predict whether AUCs for the two measurement methods will be significantly different. We fit a regression between Z scores and D values for each different NCL and used that regression to obtain a predicted value of Z for each trait pair based on D estimates for manual measurements and image-based measurements. The estimates of D for real trait pairs were 0.1 (TW), 0.14 (BN), 0.16 (TL), and 0.17 (SL), with the manually measured trait always having higher heritability than the image-based trait (Table 1). The predicted values of Z for each trait ranged from 0.15 to 0.26 when NCL=10, from 0.47 to 0.75 when NCL=100, and from 0.70 to 1.10 when NCL=1,000 (Figure 3). For each NCL we used the distribution of Z scores when D=0 and set the 2.5^th^ and 97.5^th^ percentiles as thresholds for significance to test the null hypothesis of Z=0 at α=0.05. The thresholds were (−1.26, 1.09) for NCL=10, (−1.49, 1.25) for NCL=100, and (−1.44, 1.60) for NCL=1,000. Regardless of NCL, predictions of Z for all four tassel traits fall within the thresholds for significance (Figure 3). Therefore, under the assumptions made in these simulations the manual and image-based measurements are expected to have AUCs that are not significantly different from each other.

## Discussion

In this study, we use AUCs of ROC curves constructed from GWAS results of simulated traits to test for significant differences in the ability to detect common genetic signal underlying traits with differing heritability. Our results show that as D increases, the test for differences between the traits’ AUCs becomes more significant. Though there is a strong relationship between D and Z, there is also substantial variability for Z scores at a given value of D. We predicted Z scores for real tassel morphological traits using the relationship between D and Z of simulated traits. Because each tassel trait was measured by manual and image-based methods, we predicted Z using D from the estimated heritabilities of the two measurement methods. Regardless of NCL, the predicted values of Z for real tassel traits were within the thresholds for significance that were calculated from the null distribution of Z. Based on these results, we conclude that there is unlikely to be a significant difference between AUCs of measurements made by different methods for any of the four tassel morphological phenotypes studied.

This conclusion is highly dependent on the assumptions made when creating simulated traits and performing subsequent GWAS. Here, we assumed independent and randomly positioned causal SNPs, when in fact quantitative traits can be controlled in part by numerous causal variants clustered on the same locus (Allen *et al.* 2010). Additionally, by choosing causal SNPs directly from the WiDiv-942 genotypic data, we are assuming that the causal variants are SNPs that are part of our genotypic data. The 529,018 SNPs used in this study are a small sample relative to the 60 million variants identified in the maize HapMap3 (Bukowski *et al.* 2015), and do not include other variant types that can affect quantitative traits such as insertion/deletion and copy number variants. By selecting causative SNPs from our genotyped SNPs, we make the true associations easier to find by GWAS. In reality, causative variants may not be genotyped and therefore can only be identified by linkage disequilibrium with genotyped SNPs. For simplicity’s sake, we drew the simulated effect sizes at each causal SNP from a normal distribution. Previous work by Hayes and Goddard (2001) has shown that quantitative traits in livestock appear to follow a gamma distribution with a large number of very small effect loci; they posit that there may be even more small effect loci than predicted by their distributions. This idea can be seen in its most extreme form in the omnigenic model proposed by Boyle et al. (Boyle *et al.* 2017) which is based on Fisher’s infinitesimal model (Fisher 1919). By drawing our effect sizes from a normal distribution, we may be creating more large-effect variants than is realistic, therefore increasing our ability to detect causal variants by GWAS.

Though our choices for location and effect size of causative SNPs may be increasing the probability of detecting associations, we also assume that only identifying an exact chosen causative SNP counts as a true positive. In reality, identifying associations with SNPs that are within the same gene as, or a small distance away from, the true causative SNP may be close enough. GWAS often serves as an initial sweep to find regions of interest for further study, and associations that lead to fruitful downstream analysis may still be considered a ‘success’. This is reflected in software that calculate power of GWAS by considering associations within a certain distance of the causative variant to be true positives (Liu *et al.* 2016). By only considering the exact causative SNPs as true positives we make the true positives more difficult to identify, partially counteracting the assumptions above that make the detection of associations easier.

By trying to predict the Z score for the difference between AUCs of real traits, we also make some assumptions about the tassel morphological traits that we use. Our estimates of heritability are not exact; they are population- and experiment-specific. Because the image-based traits and TW were measured in one environment, whereas TL, SL, and BN were measured in three, their estimates of heritability may have differing accuracy. By predicting Z scores for the real traits for NCL set to 10, 100, and 1,000, we were able to predict how Z scores changed as NCL changed. The true NCL for tassel morphological traits likely numbers in the hundreds or higher, with an upper limit of the total number of expressed genes (tens of thousands), as posited in the omnigenic model (Boyle *et al.* 2017). If the true NCL is greater than 1,000, the relationship between Z and D will be even steeper, meaning that the small values of D for the real tassel morphological traits may in fact be responsible for significantly different AUCs.

The use of AUC as a metric to quantify the success of GWAS is also accompanied by assumptions about the goals of GWAS. ROC curves, and thus the AUC, consider both power and type I error, as measured by true and false positive rates. Depending on the goals of the GWAS study, power and type I error may not both be of equal importance. For genomic prediction or marker assisted selection, a high type I error rate is not particularly concerning as long as power is high and trait prediction is accurate. On the other hand, studies using GWAS to choose genes for further molecular characterization have a large financial incentive to minimize type I error.

Using AUC to quantify the effectiveness of GWAS assumes that the entire ROC curve is of interest. When NCL is low this assumption may be true, but as NCL increases, it may be the case that only the beginning of the ROC curve is of practical interest. The simulated ROC curves for NCL=1,000 (Figure 2c) are close to the 1:1 line that would be achieved by randomly selecting SNPs as putatively causative. It is unlikely that a researcher would want or expect to identify every single causative locus when a trait is controlled by thousands of genes. Instead, the interest is often in large-effect loci that are likely to be identified by a stringent significance threshold. The ability to identify the most significant loci is characterized by the portion of the ROC curve close to the origin. Thus, for highly complex traits, the AUC of a partial ROC may be more informative.

In this study, we use AUC of ROC curves to characterize and quantitatively compare GWAS results from different traits. Overall, our findings show an expected relationship between NCL, heritability, and AUC of ROC curves. Greater NCL and lower heritability both reduce the AUC, while lower NCL and higher heritability can increase AUC. Results suggest that there is no significant difference between AUCs from GWAS using manual and image-based measurements of typical maize tassel characters. Creation of more nuanced simulation models and consideration of partial ROC curves may enable improvement upon the results presented in this study. The results presented here provide a foundational framework that may facilitate decision-making for researchers weighing the benefits of different phenotyping methods.

## Materials and Methods

### Plant populations: phenotyping and genotyping

The Wisconsin Diversity panel (WiDiv-942) is a set of 942 inbred maize lines that reach grain physiological maturity in the upper Midwest region of the United States (Mazaheri *et al.* 2018, in press), and represents an expanded version of the 503 line diversity panel described by Hirsch and colleagues (Hirsch *et al.* 2014). Phenotypic measurements of tassel morphology in the WiDiv-942 were performed using both manual and image-based measurements. Manual and image-based measurements are described in detail in (Gage *et al.* 2018). Briefly, manual measurements included tassel length (TL), the distance (cm) from the lowest tassel branch to the tip of the tassel spike; spike length (SL), the distance (cm) from the uppermost tassel branch to the tip of the tassel spike; branch number (BN), the total number of primary tassel branches; and tassel weight (TW), the weight (g) of the dried tassel biomass above and including the lowest branch. Image-based measurements were made using the output of the tassel phenotyping software TIPS (Gage *et al.* 2017). The data output by TIPS are image-based measurements of tassel morphology and were used as explanatory variables in a partial least squares regression model that performs image-based predictions of TL, SL, BN, and TW (referred to as TLp, SLp, BNp, and TWp, respectively) (Gage *et al.* 2018). The WiDiv-942 was grown using a replicated complete block design in three different environments: the University of Wisconsin Arlington Agricultural Research Station in the summers of 2013 and 2014, and the University of Wisconsin West Madison Agricultural Research Station in the summer of 2015. TL, SL, and BN were measured in all three environments, while TW, TLp, SLp, BNp, and TWp were measured only in 2015. Best linear unbiased predictors (BLUPs) for each inbred line for all eight traits were calculated using random effects models to account for environment, genotype-by-environment, and replication effects (Gage *et al.* 2018).

The WiDiv-942 was genotyped at 899,784 SNPs discovered by RNA sequencing (Mazaheri *et al.* 2018, in press). SNP data contained 30% missing data, which were imputed using fastPHASE (Scheet and Stephens 2006). After imputation, 0.3% of SNP calls were missing, due to inability of the imputation program to call all missing SNPs. The remaining missing data at any given SNP were imputed to the mean value for that SNP. SNPs with a minor allele frequency of <0.02 were removed from the genotypic data, leaving 529,018 remaining SNPs.

### Simulated phenotypes

Heritability (*h*^2^) is the ratio of genetic variability 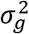 in a population to overall phenotypic variability 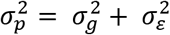, where 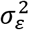 is all variance not attributed to differences between genotypes. Thus, 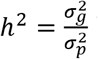is a measure of the strength of genetic signal in a particular population for a particular phenotype. Two different measures of the same character, for instance, TL and TLp, can be thought of as correlated traits with the same underlying genetic control but differing heritabilities. The true value of the character for a particular individual cannot be measured perfectly, but both the manual and image-based measurements of the character represent a combination of true signal and some (differing) amount of measurement error.

To simulate traits with similar behavior, effect sizes were randomly drawn from a normal distribution for a set of randomly chosen SNPs genotyped in the WiDiv-942, and the ‘true’ phenotypic value was calculated for each of the 942 individuals. In a second step, noise was added to each individual’s true value in order to attain a desired heritability. We varied both the number of causative loci (NCL), to simulate traits controlled by differing numbers of variants, and the heritability of simulated traits. The NCL was set to 10, 100, and 1000, and *h*^2^ ranged from 0.1 to 0.9 in increments of 0.1. The causative loci were randomly selected a single time from all SNPs genotyped in the WiDiv-942. The phenotype for an individual *i* for a trait controlled by NCL=*n* SNPs and heritability *h*^2^ is 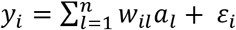. The standardized genotypic value 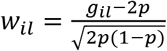is the genotype *g_il_* of individual *i* at SNP *l*, expressed as 0, 1, or 2 copies of the major allele, centered by twice the major allele frequency, *p*, and divided by the standard deviation of the SNP. The allelic effect *a_l_* is drawn from *N*(0, 10), and *ε_i_* from 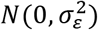 where 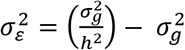 The variance for allelic effects was set arbitrarily, but could be any reasonable number as the error variance is modified to ensure the desired heritability. Genotypic variance 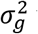 was calculated simply as the sample variance of the population’s true phenotypic values. For each combination of *n* and *h*^2^, phenotypes were simulated 10 times. The difference between each simulation with the same set of parameters is simply the *>_i_* representing random draws from the same distribution each time. In total, 270 traits were simulated: pairwise combinations of _3_ levels of *n* and 9 levels of *h*^2^, each replicated 10 times.

### Genome-wide association studies

Genome-wide association studies (GWAS) were conducted with the software GAPIT (Lipka *et al.* 2012), implemented in R (R Core Team 2016). The same kinship matrix, calculated from 10,000 randomly selected SNPs by the VanRaden method in GAPIT (VanRaden 2008; Lipka *et al.* 2012), was used for all GWAS and no other covariates (e.g., principal component scores) were included in the model. Compression was turned off by setting the group.to and group.from parameters to 9999. GWAS was run separately for each of the 270 simulated traits.

### Receiver operator characteristic curves

Receiver operator characteristic (ROC) curves are typically used to assess the ability of a particular method for identifying the true, binary status (case or control) of an individual based on various threshold levels of a continuous predictor variable. The *roc()* function in the R package pROC (Robin *et al.* 2013) was used to construct ROC curves from GWAS results by considering each SNP as an ‘individual’ and coding the randomly selected causative SNPs as cases in the response, while all other SNPs were considered controls. The −log_10_(p-value) of each SNP was used as the predictor variable. A separate empirical ROC curve was fitted to the GWAS results for each of the simulated traits. A representative ROC curve was estimated from all 10 replicates of each parameter combination by calculating the mean of the sensitivities and specificities along the curve.

The area under the curve (AUC) for an empirical ROC curve is equivalent to a Mann-Whitney statistic calculated on the predictor scores of the case and control individuals (DeLong *et al.* 1988). In addition, the AUC can be interpreted as the probability that for any pair of randomly selected case and control individuals (or in this case, causative and non-causative SNPs), the predictor value of the control individual will be less than or equal to the predictor value of the case individual (DeLong *et al.* 1988). In this context, that is the probability that the −log_10_(p-value) of the non-causative SNP is less than or equal to the −log_10_(p-value) of the causative SNP. Because the empirical AUC of an ROC curve acts as a Mann-Whitney statistic, the difference between AUCs of two ROC curves can be nonparametrically tested (DeLong *et al.* 1988), yielding a statistic we refer to as Z. Because our Z scores were not standard-normally distributed as expected (DeLong *et al.* 1988), we instead used the values of Z when the difference in heritability (D) between traits was 0 as a null distribution for a given NCL. Thresholds for significance at α=0.05 were calculated empirically from the 2.5^th^ and 97.5^th^ percentiles of the null distribution of Z.

Z-scores were calculated for tests of the difference between all pairwise combinations of ROC curves with the same NCL using the *roc.test()* function in pROC with the method argument set to ‘delong’ (Robin *et al.* 2013).

### Estimating Z scores and confidence intervals for real phenotypes

Because the relationship between Z and D for pairs of simulated traits was approximately linear, we fit a simple linear regression of Z against D separately for each value of NCL. Using the differences between estimated heritability for each pair of real traits (TL and TLp; SL and SLp; BN and BNp; TW and TWp), we predicted the value of Z for each character at different NCL using the fitted regressions. Using the thresholds for significance derived from the empirical distribution of Z when D=0, we tested the null hypothesis Z=0 for each character at each NCL.

## Data Availability

Genotypic data are available from doi:10.5061/dryad.n0m260p (paper still under review; doi not publicly available as of submission). Scripts for analysis can be found at github.com/joegage/GWAS_AUC

## Acknowledgements

### Funding

JG is supported by the National Research Initiative for Agriculture and Food Research Initiative Competitive Grants Program grant no. # 2012-67013-19460 from the USDA National Institute of Food and Agriculture. This work was also supported by the USDA National Institute of Food and Agriculture, Hatch projects WIS01923 and WIS02020.

### Authors’ contributions

All authors contributed ideas, and all authors were involved in writing, editing, and approving the final manuscript.

